# Overexpression of miR5505 enhanced drought and salt resistance in rice (*Orayza sativa*)

**DOI:** 10.1101/2022.01.13.476146

**Authors:** Yuanwei Fan, Jiankun Xie, Fantao Zhang

## Abstract

Rice is one of the most important crops in the world and half of the world population consumes it as their staple food. The abiotic stresses caused by drought, salt and other stresses have severely impacted rice production. MicroRNAs (miRNAs) are a type of small non-coding RNAs which widely reported as gene regulators, suppressing genes expression by degradation mRNA or translation inhibition. Previously, high-throughput sequencing has found a conserved miRNA miR5505 responding to drought stress in Dongxiang wild rice (DXWR). Several other studies also revealed that miR5505 was involved in rice stress responses. We further studied the effect of miRNA in drought and salt tolerance by overexpression it in rice. 2 in 18 successfully transformed transgenic lines with higher miR5505 expression were selected and then drought and salt resistance ability were evaluated. Both transgenic lines showed stronger drought and salt tolerance than wild-type (WT). Putative targets of miR5505 were identified by psRNATarget and several of them were found stress-related. RNA-seq found 1,980 differentially expressed genes (DEGs) in transgenic lines. Among them, 978 genes were down-regulated. Three genes were predicted by psRNATarget and two of them might be stress-related. We also found various environmental stress *cis*-acting elements in upstream of miR5505 promoter through Software PlantCARE. In all, we improved rice drought and salt tolerance by overexpressing miR5505, and the generated putative targets and *cis*-acting elements also suggested miR5505 might play important roles in the regulation of drought and salt responses.

## Introduction

Rice is one of the leading crops in the world. Half of the world’s population consumes it as their staple food^[1]^. As a water-intensive crop, rice, unfortunately, makes its production challenged when encountering unexpected drought. While global temperature keeps rising caused by climate change, it accelerates evaporation which makes arable land suffer more drought and become saltier. It has been estimated that one degree of temperature increase will result in tropical rice production decreasing by 1.3%-3.5%^[2]^. Temperature rising might cause more unexpected extreme weather^[3]^, which will further reduce rice production^[4]^. In Southeast Asia, about 23 million hectares of water-fed rice have been significantly affected by the unexpected drought^[5]^. Securing its production mainly relies on arable soil and the abundance of water, while more than 10% of arable land is affected by drought and salinity^[6]^. Most widely cultivated rice shows high drought and salt intolerance, and the abiotic stress caused by salinity greatly affected rice yield across the world^[7]^. Thus, rice breeding for stronger environmental adaptation to abiotic stresses like drought and salt stress is crucial and urgent.

In plants, microRNAs (miRNAs), which have a length of 21-24 nucleotides (nt), involve in various developmental processes, including biotic and abiotic defenses^[8–10]^. The mechanism of the miRNA mediating gene expression is mainly through repressing the gene by sequence complementation with target transcripts, which will lead to degradation or translation inhibition to those targets^[11]^. To date, many miRNAs have been proved that they play important roles in abiotic stress adaptation, such as drought^[12]^, salt^[13]^, cold^[14]^ and metal, which include aluminum^[15]^, arsenic^[16]^, Cd ^[17]^ and so on. High-throughput sequencing technology has helped us better understand the profile of miRNAs in mediating drought stress in rice^[18–20]^, and has identified a group of miRNAs that play important roles in rice drought defense^[21]^.

Dongxiang wild rice (Oryza rufipogon, DXWR) is a common wild rice that is discovered in Dongxiang county, Jiangxi province. It is the northernmost common rice that we have found in the world^[22]^. Because it can resist to various abiotic stresses^[23, 24]^, DXWR is considered a very precious rice genetic resource. We recently identified SSR markers, especially drought-related SSR markers and storability-related loci from DXWR ^[54, 55, 56]^. Previously, by high-throughput sequencing technology, we have identified several miRNAs which might be related to drought stress responses in Dongxiang wild rice (DXWR, *Oryza rufipogon* Griff.)^[25]^. Among those miRNAs, miR5505 was selected for further research. Researchers have found that the DNA methylation of miR5505-generating loci was higher in rice when exposed to cadmium^[26]^. It has been widely reported that the global genome methylation occurred when tissue exposed to the abiotic stress^[27]^. Expression of miR5505 is higher in heat stress tolerance rice strain than the susceptible ones. After high-temperature treatment for 24h, the difference got larger ^[28]^. However, further research of miR5505 in regulating abiotic stress response in rice is still unavailable. Here, we overexpressed miR5505 and found improved drought and salt resistance in transgenic lines, suggesting that miR5505 might have a positive effect on drought and salt stress resistance. To investigate miR5505 regulatory pathway, psRNATarget and PlantCARE were deployed to predict putative targets and *cis-acting* elements, respectively. We also investigated the global gene expression profile in miR5505 transgenic lines, and found 1,980 differentially expressed genes (DEGs). The result suggested that miR5505 might be a very important miRNA in rice drought and salt response regulation family.

## Materials and Methods

### Plant materials, drought and salt stress treatment

The rice plants that were used in this study included Dongxiang wild rice (DXWR, *Oryza rufipogon* Griff.), cultivated rice Zhonghua 11 (ZH11, *Orayza sativa* L. subsp. *japonica*) and miR5505 transgenic lines (ZH11 background). Rice seedlings were cultured at 26 ± 2 ^o^C with a relative humidity of 75 ± 5% for 14 h light/10 h darkness. Hydroponic culture system with IRRI nutrient solution was applied to cultivate those plants as described^[29]^. At the stage of four leaves, seedlings’ nutrient solution was poured out for drought treatment, and nutrient solution supplemented with 200 mM sodium chloride for salt treatment, respectively. For miR5505 expression analysis, seedlings of miR5505 overexpression lines were collected at four leaves stage. All experiments were repeated at least three times.

### Vector construction and transformation

To generate miR5505 overexpression transgenic lines, precursor miR5505 (pre-miR5505) sequence including 699 bp upstream and 252 bp downstream sequence of miR5505 was pulled out from DXWR genomic DNA with primer OsmiR5505-F and OsmiR5505-F (supplementary table1). To construct the miR5505 overexpression vector, the PCR product of pre-miR5505 was cloned into the *KpnI* and *SalI* restriction sites of the binary vector pCAMBIA1300 which carries the hygromycin phosphotransferase gene (*hph*) as the selectable marker. After confirmation of the correct insertion by sequencing, the constructed vector was transformed into ZH11 seedling by *Agrobacterium tumefaciens-mediated* method^[30]^.

### RNA extraction, qRT-PCR, psRNATarget prediction, RNA-seq and cis-acting elements analysis

Total RNA was extracted by TRIzol (Sangon Biotech, China) as the manufacturer’s manual described. Then, 3-4 ng RNA was reverse transcribed using miRNA First-Strand cDNA Kit (Sangon Biotech, China). qRT-PCR was carried out by using TB Green Premix Ex Taq II (Tli RNaseH Plus) kit (Takara, China) on ABI7500 Real-time system (Applied Biosystems, USA). Rice *U6* snRNA gene was used as the control for miRNA expression analysis. 2^-ΔΔCt^ method was applied for data analysis. All assays were carried out in triplicate under identical conditions. All primers used in this study were listed in the supplementary table1. Target prediction software100 psRNATarget was used to predict miRNA5505 targets with default settings of Schema V2 (2017 release)^[31]^. The cDNA library reference is MSU Rice Genome Annotation (version 7). RNA-seq and the subsequent analysis were carried out by biomarker (Shandong, China). The reference genome database came from Plantbiology (MSU_v7.0). Software PlantCARE^[32]^ was used to identify Putative cis-acting elements in the miR5505 promoter which includes ~1500 bp DNA upstream sequence of miR5505 in the DXWR genomic DNA.

## Results and Discussion

### Overexpression of miR5505 enhance drought tolerance

The overexpression of miRNAs has been widely used to study the roles of miRNA in abiotic stress tolerance response, such as miR164^[33]^, miR319^[34]^, and miR408^[35]^. To investigate the role of miR5505 in rice in response to drought and salt stress, miR5505 overexpression transgenic lines were generated. PCR test amplified by primers for *hph* was used to identify positive transgenic rice lines after screening for hygromycin resistance. Further studies were carried out on T3 transgenic homozygous rice line. Out of 18 independent transgenic lines, two lines that showed relatively high accumulation of mature miR5505 were chosen for further studies (named miR5505OE-1, miR5505OE-2) (Fig. 1). The two transgenic lines do not show any abnormal phenotypes compared with wild-type (WT) in the seedling growth stage (Fig. 2A, D). At four leaves stage, after pouring out culture solution to subject water deprivation for 24 hours, both WT and transgenic lines were wrinkled. Then the seedlings were re-watered with culture solution for 7 days to recover, both WT and miR5505 overexpression lines were partially recovered (Fig. 2B, E), while the transgenic lines grew better than WT. The survival rate of WT and miR5505OE-1 were 50.0% and 91.4%, respectively, and showed significant differences (Fig. 2C), while WT and miR5505OE-2 were 64.8% and 88.3%, respectively, and showed significant differences, too (Fig. 2F). The results suggested that overexpression of miR5505 increased rice drought tolerance.

**Fig. 1.**
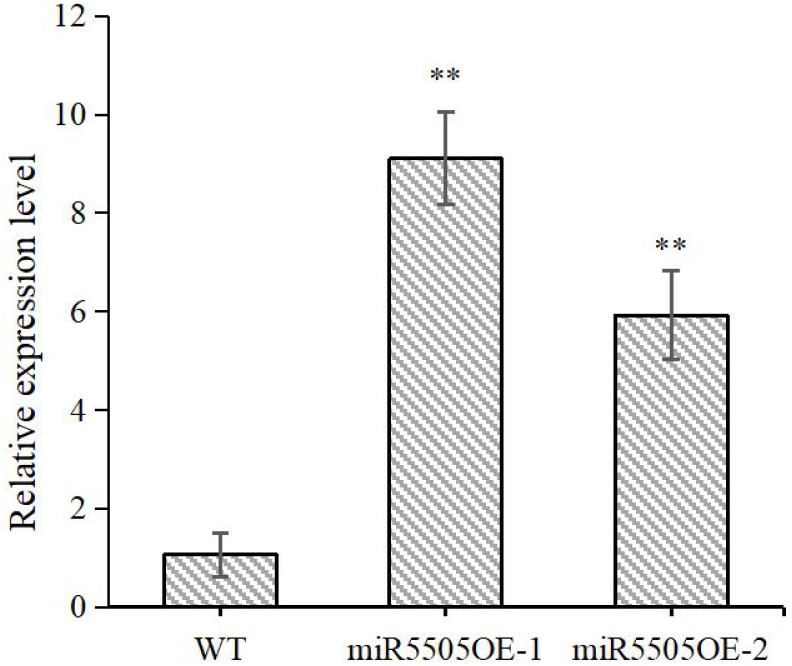
miR5505 relative expression in WT and transgenic lines in the normal conditions. Error bars show the standard deviations ofmean, **represents significant differences at *P* < 0.05 according to Student’s *t*-tests.

**Fig. 2.**
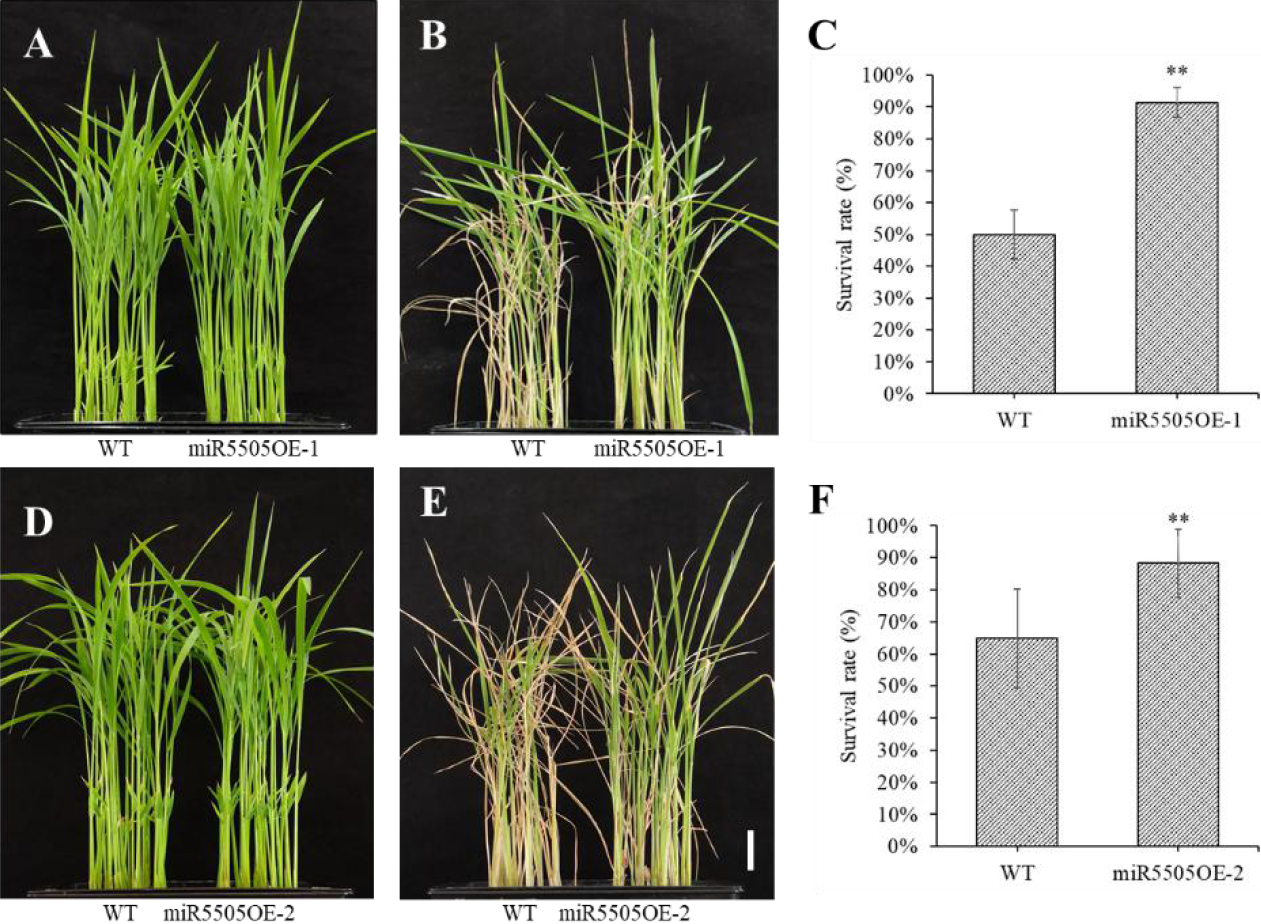
Drought tolerance evaluation of miR5505 transgenic rice. Seedlings grew to four leaves stage and then the nutrient solution was poured out to subject water deprivation for 24 hours. (A, D) WT and transgenic rice seedlings before treatment, (B, E) recovery for 7 days and (C, F). The survival rate of WT and transgenic lines after recovery. Scale bar = 3 cm, Error bars show the standard deviations of mean, ** represent significant differences at *P* < 0.05 according to Student’s *t*-tests.

The global mean temperature caused by climate change steadily increases, which prolongs the dry seasons and makes the drought intensity^[36]^. Up to $30 billion of crop production was lost caused by drought in the last decade^[37]^. Rice is generally grown in water-sufficient condition which makes it severely susceptible to water deprivation. So, accelerating rice breeding with stronger drought tolerance is important and urgent. To facilitate the process, research of miRNA could help us better understand rice drought-adaptive mechanisms and eventually help rice breeding. It was widely reported that miRNAs overexpression enhanced drought tolerance of the transgenic plants, but the result varies when the same miRNA overexpressed in different species^[38]^. Here, we overexpressed miR5505 and found transgenic rice drought tolerance ability improved. The mechanisms of miR5505 regulation needs to be carefully studied for the sake of rice breeding with drought tolerance.

### Overexpression of miR5505 enhances salinity tolerance

We also investigated the salt tolerance of miR5505OE-1 and miR5505OE-2 transgenic lines. We transferred them to the nutrient solution supplemented with 200 mM NaCl and treated them for five days. Most of the leaves of transgenic lines were almost normal and some were still slowly growing, while most leaves of WT were wilted and stopped growing. After being transferred to the nutrient solution without salt and continued to cultivate for another 5 days, both miR5505OE-1 and miR5505OE-2 were mostly recovered, while WT were severely affected by salt stress and most of the specimens died (Fig. 3B, E). This result reflected that transgenic lines of miR5505OE-1 and miR5505OE-2 had higher survival rate (WT and miR5505OE-1 were 28.1% and 78.1%, respectively; WT and miR5505OE-1 were 25.0% and 75.0%, respectively) (Fig. 3C, F). The results suggested that miR5505 transgenic lines had higher salt tolerance.

**Fig. 3.**
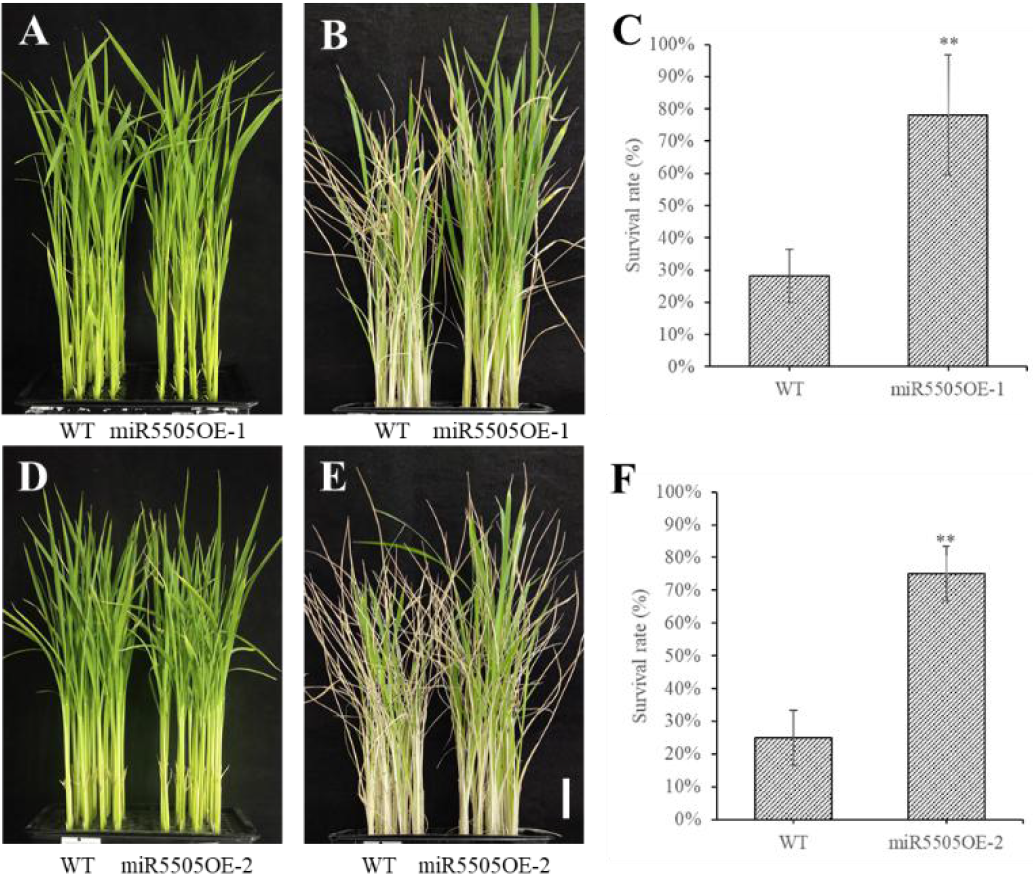
Salt tolerance evaluation of miR5505 transgenic rice. Seedlings grew to four leaves stage and were treated with the nutrient solution supplemented with 20 mM sodium chloride for 5 days. (A, D) WT and transgenic rice seedlings before treatment, (B, E) recovery for 5 days and (C, F). The survival rate of WT and transgenic lines treated with 20 mM sodium chloride. Scale bar = 3 cm. Error bars show the standard deviations of mean. ** represents significant differences at *P* < 0.05 according to Student’s *t*-tests.

After the drought, salinity is the second soil problem in the countries of rice planting and is believed to be a big obstacle to rice yield increase worldwide^[39]^. High soil salinity has negatively affected one-fifty of irrigated agriculture. Rice, which feeds half of the world population and is the most salt-sensitive cereal^[40]^, is badly needed to improve salinity tolerance in the future. miRNA has been widely reported involve in salinity stress response^[41, 42]^. Some miRNAs expression changes could alter rice salinity^[43, 44]^. We find miR5505 overexpression has improved the rice salt tolerance, which makes it a meaningful candidate miRNA in the future to study salt resistance of rice. To date, seldom studies about miR5505 has been reported in the area of the abiotic stress responses. More researches have to be done to elaborate the full map of the miRNA stress tolerance mechanisms.

### Targets prediction and analyze *cis*-acting elements of miR5505

miRNAs suppress target genes’ expression through sequence-specific cleavage or translational inhibitioní^[11]^. To probe the regulatory targets of miR5505, psRNATarget software was deployed and 116 putative candidates were found (supplementary table 2). Among them, 100 were labeled as “cleavage” and the rest were “translation”. 25 of the putative targets were described as retrotransposon, which was widely reported involve in plant stress responses^[45]^.9 putative targets’ description was “disease resistance”. Those biotic stress genes could also involve in abiotic stress responses^[46]^. 4 putative targets encoded lipases which had been reported involve in plant abiotic stress responses^[47]^ (supplementary table 2).

As a regulator, its gene expressions are also regulated by its upstream regulators. Motifs in the promoter region play key roles in upstream regulator’s recognizing and binding. Transcription factors (TFs) change plant behaviors by altering gene expression through binding its special DNA sequence in the genes’ upstream promoter region. Various *cis*-acting elements in the promoter region may lurk regulatory information of the miRNA gene. Numerous studies have shown that motifs play important roles in abiotic stress responses^[48]^. About 1.5 kb DXWR genomic sequence upstream of miR5505 was pulled out and blasted against PlantCARE database to search for *cis*-acting elements. 12 potential *cis*-acting elements related to environmental stresses were identified in the upstream of miR5505 gene. One *cis*-acting element is involved in the abscisic acid (ABA) responses. ABA is a phytohormone that is critical for plant growth and integrates various stress signals to control subsequent stress response^[49]^. This suggests that miR5505 regulating abiotic stress might be affected by the ABA signal pathway. We also found “MBS” which is an MYB binding site is involved in drought inducibility. This might explain why miR5505 transgenic rice has higher drought tolerance.

Therefore, fully understanding upstream *cis*-acting elements and putative targets of miR5505 might reveal the mechanisms of how the miRNA regulates abiotic stresses, which further help genetic engineering and other approaches to improve rice drought and salinity resistance.

**Table 1.**
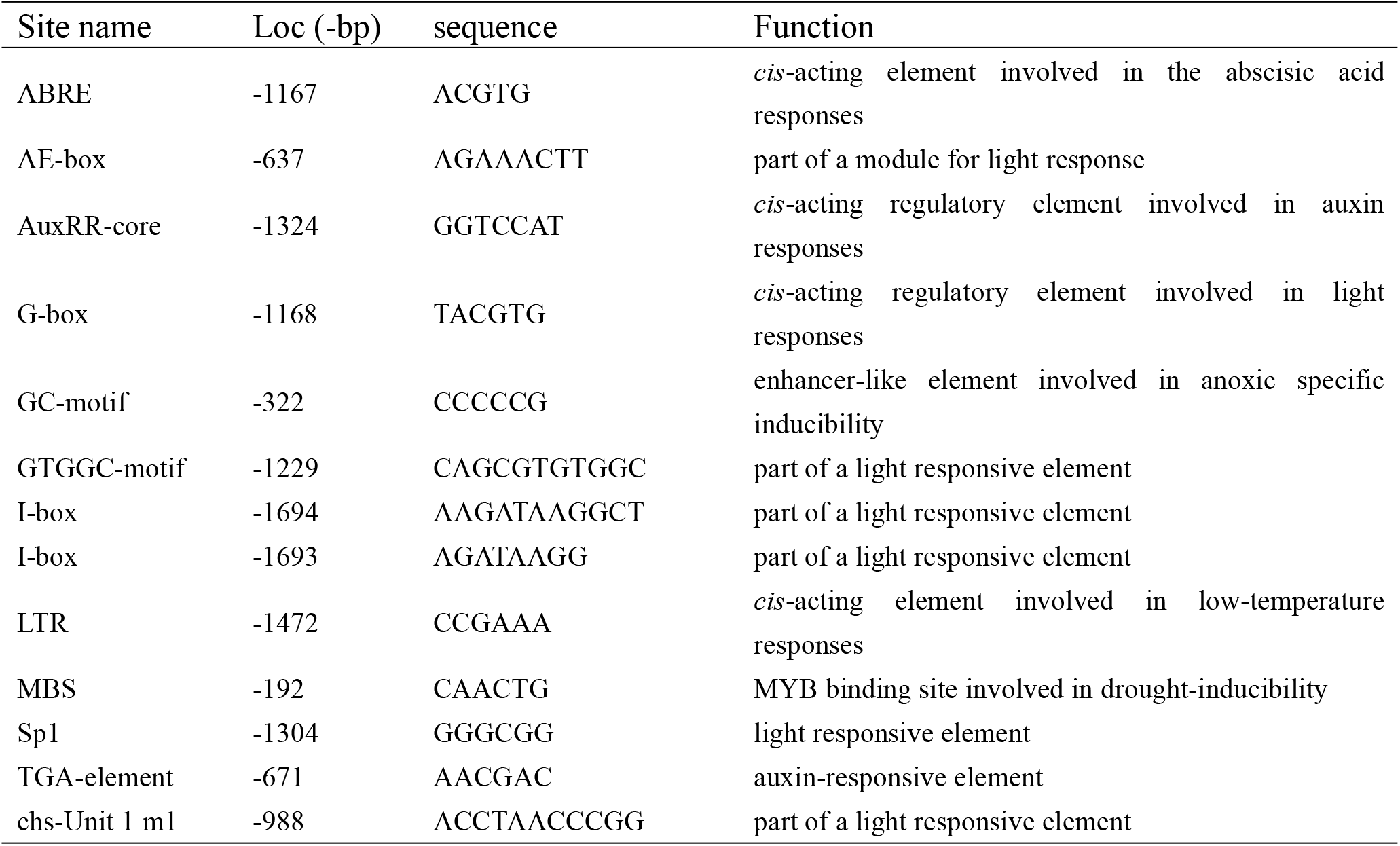
List of stress-related *cis*-acting elements in upstream of miR5505 promoter

### Overexpression of miR5505 results in global gene expression changes in rice

To investigate how the putative targets of miR5505 respond to drought and salt stresses after miR5505 overexpression, RNA-seq assay of wild-type and overexpression transgenic lines were carried out in normal conditions (Fig. 4A). The average total-mapped ratio of the reference genome was 95.06%, while among them, the unique-mapped ratio was 95.80%, with an average of 49,820,884 reads mapped with the reference genome. Differentially expressed genes (DEGs) are defined as the genes that the fold changes are more than two times and false discovery rates (FDR) are less than 0.01. Compared with WT, a total of 1,980 DEGs were found in miR5505 transgenic lines. Among them, 978 genes were down-regulated, while 1,002 genes were up-regulated (Fig. 4B). Gene ontology (GO) function classification analysis was further employed to figure out the function of these DEGs (Fig. 4C). A total of 611 significant DEGs were related to metabolic processes, while 426 were related to cellular processes, suggesting that miR5505 affected biological processes in transgenic lines. In the GO term of cellular component, membrane (471 DEGs) and membrane part (429 DEGs) were significantly enriched, also suggesting that miR5505 affected membrane and cellular parts in transgenic lines. Further, binding (666 DEGs) and catalytic activity (720 DEGs) also revealed that miR5505 affected molecular function in transgenic lines (supplementary table 3). All those results suggest that miR5505 overexpression in rice has globally altered the gene expression and could affect its ability of drought and salt tolerance.

**Fig. 4.**
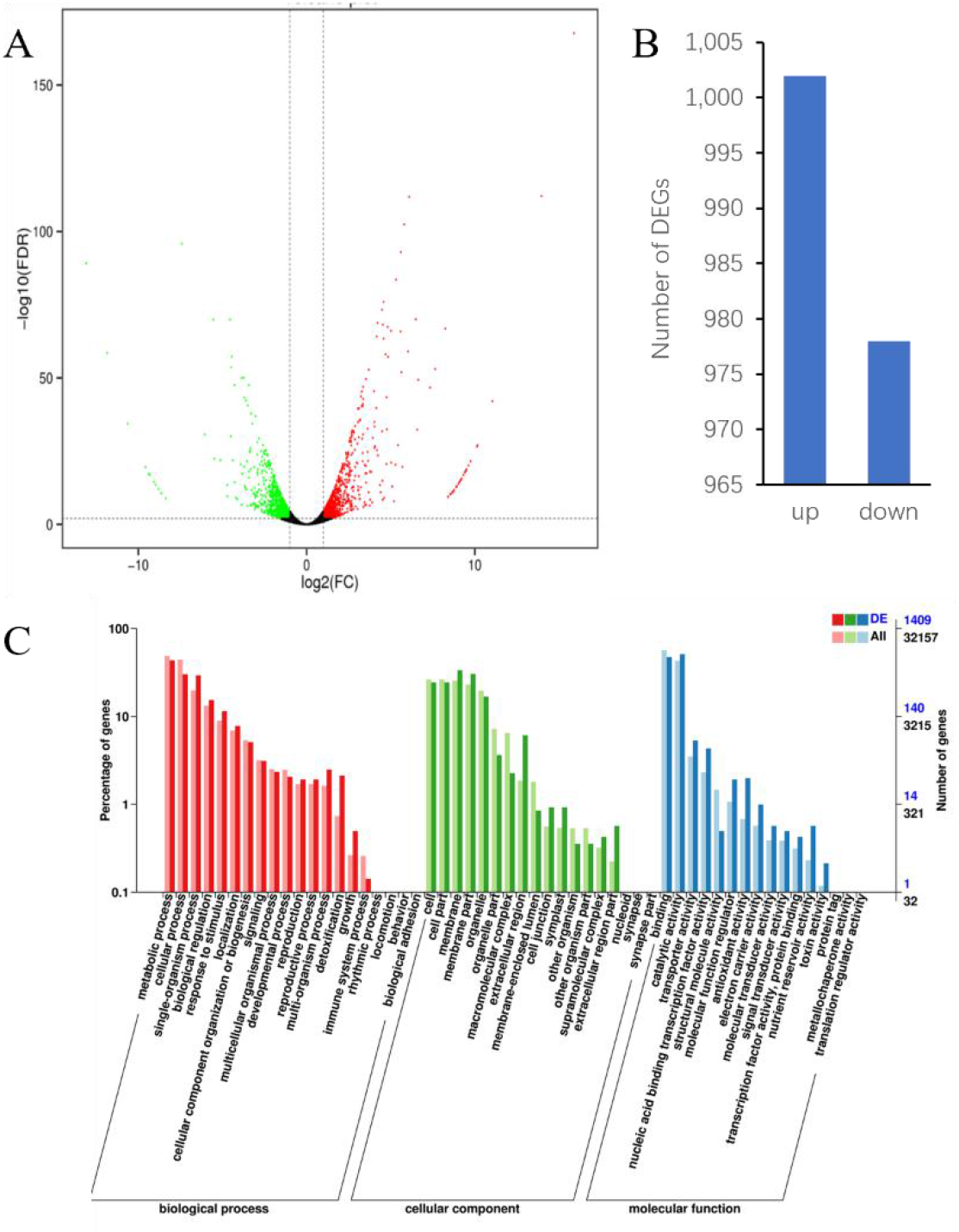
Overexpression of miR5505 resulted in global gene expression changes in rice. (A) volcano plot of DEGs identified by RNA-seq in WT and miR5505 transgenic lines. (B) clustering analysis of DEGs in WT and miR5505 transgenic lines, and (C) gene ontology (GO) classification for the identified target genes.

RNA-seq results could reveal gene expression profiles at a large scale, which can be used to investigate the differential gene expression in various tissues or specimens. Based on RNA-seq data, various pathways could be revealed that are involved in miR5505 regulated drought and salt stress responses in transgenic lines. Studies showed that hormone signaling pathways involved in biotic stress could also be a participant in abiotic stress pathways^[50]^. Here we found 97 DEGs in miR5505 transgenic lines involved in plant-pathogen interaction, which might also be abiotic stress-related (supplementary table4). Phenylpropanoid biosynthesis pathway is active when encountering abiotic stresses like drought and salinity, which will lead to the accumulation of phenolic compounds that play an important role in the physiological and metabolic processes^[51]^. Here, we found 59 DEGs expressions in phenylpropanoid biosynthesis pathway in miR5505 transgenic lines (supplementary table4), suggesting elevated miR5505 expression boosted abiotic related metabolic process to improve abiotic stress tolerance in rice.

The function of miRNA repressing the gene expression is mainly by sequence complementation with target transcripts, which leads to degradation or translation inhibition to those targets^[11]^. Here the expression level of three DEGs (*LOC_Os09g18230, LOC_Os04g22910, LOC_Os04g39610*), which were predicted by psRNATarget, were decreased in the miR5505 overexpression lines. *LOC_Os09g18230* belongs to the receptor-like cytoplasmic kinase (OsRLCK) gene family. Some RLKs have been characterized and are found to play roles in plant development and stress responses^[52]^. *LOC_Os04g39610* encodes a putative glycerophosphoryl diester phosphodiesterase, which might be involved in lipid metabolisms induced by abiotic stress^[53]^.

miRNAs responses to stress also depends on the functions of its targets. The three targets are significantly down-regulated in the miR5505 overexpression plants, suggesting miR5505 might improve drought and salt tolerance through them. Further research should be implemented to expatiate the function of the target genes and investigate the cleavage mechanisms between miR5505 and those targets.

## Conclusion

In general, our results showed that overexpression of miR5505 in rice enhanced plant drought and salt tolerance without changes its phenotypes in optimal conditions. Further research should be done to understand miR5505 regulation mechanisms of abiotic stress tolerance, which could provide clues to understand the miRNAs in response to abiotic stress, and ultimately help high-stress tolerance rice breeding.

## Acknowledgment

The sponsor of this research was Jiangxi Provincial Key Lab of Protection and Utilization of Subtropical Plant Resources, College of Life Sciences, Jiangxi Normal University, Nanchang 330022, China. (YRD201913), Postgraduate Innovation Fund of Jiangxi Normal University, (YC2019-B043).

